# Harnessing Circadian Resilience for Demand-Flexible Controlled Environment Agriculture

**DOI:** 10.64898/2026.11.12.765898

**Authors:** Martin William Battle, Zoë Grace McFarlane, Csaba Hornyik, Philip Prizeman-Green, Tanveer Khan, Jethro Browell, Matthew Alan Jones

**Author notes:** Author for correspondence: The author responsible for correspondence is Prof. Matt Jones.

## Abstract

The increasing penetration of renewable electricity generation creates opportunities for flexible demand-side energy management in Total Controlled Environment Agriculture (TCEA), where artificial lighting represents a major operational cost. However, the biological consequences of demand-responsive lighting strategies remain poorly understood. Here, we investigated whether photoperiods can be dynamically adjusted to minimise economic costs or maximise renewable energy utilisation while maintaining crop productivity. Using historical electricity market and wind-generation data, we evaluated three demand-flexing strategies: short-term demand-side response (DSR) events, cost-optimised photoperiods aligned with low electricity prices, and wind-optimised photoperiods aligned with periods of high renewable generation. Despite substantial variation in photoperiod length, plants maintained robust circadian rhythms under both cost- and wind-optimised regimes, demonstrating considerable resilience of the circadian system to irregular environmental timing. Commercial-scale trials involving basil, lettuce, mustard greens and rocket revealed that demand-flexed lighting regimes had minimal effects on harvested fresh or dry weight. Our findings demonstrate that TCEA can provide meaningful demand-side flexibility and support renewable energy integration without substantial reductions in crop productivity. Circadian resilience appears to buffer plants against fluctuating photoperiods, providing a biological foundation for the deployment of demand-flexible lighting strategies in future low-carbon agricultural systems.

## Introduction

Total Controlled Environment Agriculture (TCEA) offers a transformative approach to crop production by enabling year-round, high-quality yields independent of the effects of changing seasons or climate change (Kaiser et al., 2024). This approach is particularly valuable in regions where short growing seasons, extreme temperature, suboptimal soils, and inconsistent daylight limit conventional outdoor farming. Although the benefits of TCEA require a significant energy investment, the rapid, worldwide expansion of renewable electricity generation offers a potential solution for growers exposed to increasingly volatile energy markets (de Carbonnel et al., 2022; Kaiser et al., 2024). Evolving smart energy grids benefit from flexible electricity consumption since renewable energy sources have more varied generation than fossil-fuelled alternatives (Khalid, 2024). TCEA facilities could therefore provide both economic returns and system-level benefits by serving as energy management tools in addition to producing crops.

TCEA can implement end use energy demand via several strategies. In its simplest form, commercial Demand Side Response (DSR) contracts allow the grid operator to curtail energy consumption for a limited period with limited notice (2.5-5 h, NESO 2026). Alternatively, TCEA facilities could strategically adjust their electricity consumption in response to real-time grid conditions and pricing signals (Khalid, 2024). By aligning electricity use with periods of high renewable energy availability or low market prices, energy-intensive sectors such as TCEA can reduce costs while contributing to grid stability and decarbonisation.

The optimisation of lighting regimes must balance economic and environmental benefits against potential impacts on crop performance, such as yield, quality, and uniformity—traits that are critical for the commercial viability of TCEA (Kaiser et al., 2024). Lighting alone can account for up to 85% of energy use in indoor farming, making it a primary target for optimisation (de Carbonnel et al., 2022; Kaiser et al., 2024). However, variations in light timing and intensity can alter plant development, introduce variability between individuals, and complicate harvesting processes (Liebers and Pfannschmidt, 2024). Understanding these trade-offs is consequentially essential for the successful deployment of demand-responsive TCEA systems.

Plant growth is strongly influenced by light availability, with a complex network of photoreceptors and signalling pathways coordinated by the circadian clock to align environmental responses and developmental transitions with prevailing conditions (Battle et al., 2024; Jones, 2018; Sanchez et al., 2020). This sophisticated sensory system confers a substantial adaptive advantage in heterogeneous and resource-limited natural environments, enabling plants to optimise growth and resource acquisition. However, such responsiveness can present challenges for intensive agricultural systems, where uniformity in yield, quality, and harvest timing is desirable to support mechanised production and reliable commercial supply chains (Kaiser et al., 2024; Li et al., 2018; Palmer et al., 2012).

In TCEA systems, the circadian clock has emerged as a potential target for improving crop performance, sometimes described as ‘chronoculture’ (Steed et al. 2021). Environmental interventions are timed to coincide with specific phases of the circadian cycle to maximise beneficial physiological responses and enhance yield or quality (Steed et al., 2021). However, the implementation of chronoculture must be balanced against the phenotypic plasticity of plants, which can dynamically adjust circadian and environmental sensitivity in response to changing growth conditions.

Understanding how circadian regulation interacts with environmental adaptation is essential for determining the robustness and transferability of time-of-day-based cultivation strategies into TCEA. Phenotypic plasticity can be restricted by over-expression of a constitutively-active allele of phytochrome B (Y276H, YHB) or by disruption of the circadian system (Battle et al., 2024). However, it remains unclear whether the benefits of consistent performance outweigh the loss of circadian regulation of growth and development.

In this study, we evaluate the potential of end user energy demand management in TCEA. We assess how adaptive lighting regimes can be aligned with real-time grid conditions to minimise economic or environmental costs while maintaining crop productivity. We also examine the extent to which genetic and physiological approaches can reduce phenotypic plasticity to maintain yield under variable lighting. Through a combination of experimental trials and commercial-scale demonstrations, we identify operational thresholds at which application of variable photoperiods delivers meaningful reductions in energy cost and carbon emissions without compromising agricultural performance.

## Materials and Methods

### Plant material and growth conditions

Wild-type *CCA1::LUC+* and *elf3-2 CCA1::LUC+ Arabidopsis* seeds have been reported previously (Huang et al., 2016). *PHYB::YHB* and *PHYB::YHB* (*elf3-2*) *Arabidopsis* have been reported previously (Battle et al., 2024). Basil (Sweet Genovese), Lettuce (Green Batavia, Tango), Mustard (Green Frills), Salad Rocket (Victoria) and Watercress seed were all purchased from CN Seeds (Cambridgeshire, United Kingdom).

### Luciferase assays

Seeds were surface sterilized in chlorine gas and stratified in sterile water at 4°C for at least 3 days prior to plating individually on half-strength Murashige and Skoog (0.5 MS) medium (Prasetyaningrum et al., 2023). Seedlings were grown for 12 days under 50 μmol m^−2^ s^−1^ white LED light in the corresponding experimental light regime shown in the figure legends in LMS cooled incubators with temperature was set to 22°C.

Around 18 h before imaging began, plants were sprayed with 3 mM D-luciferin in 0.1% Triton X-100 and returned to entrainment conditions. Individual plants were imaged repeatedly (every 2 h), using a Retiga LUMO camera run by MicroManager 1.4.23 using a custom script (Edelstein et al., 2014). Temperature was maintained at 22°C throughout imaging. Plants were imaged under 50 μmol m^−2^ s^−1^ white LED light with the same experimental light regime used during the growth of the given plants. The patterns of the luciferase signal were fitted to cosine waves using Fourier fast transform non-linear least squares (Plautz et al., 1997) to estimate the circadian period length made using BioDare2 (Zieliński et al., 2022 ; www.biodare2.ed.ac.uk). Relative amplitude error (RAE) was calculated by dividing the amplitude error estimate for each curve by the amplitude value (Plautz et al., 1997). Data were considered rhythmic if the fitted curve returned a period estimate within 16–30 h and had an RAE < 0.5. Waveforms, periods, and percentage rhythmicity data were plotted using GraphPad Prism v.11.0.2.

### Experimental light regimes

For growth experiments, plants were grown under one of four light regimes with different timings of light and dark periods but the same light qualities and intensities; Control, Demand Side Response (DSR), Cost-optimised, or Wind-optimised. Control conditions comprised long days (18 h light: 6 h dark cycles). DSR plants were grown under long days (18 h light: 6 h dark cycles) with a 15-minute dark interval every 4 h, beginning from 1 hour after dawn (Figure 1A). Cost-optimised plants were grown under a variable cycle where 6 h dark periods were aligned with the most expensive continuous 6 h period for energy according with energy grid data (Figure 1A, 1C). Similarly, wind-optimised plants were grown under a variable cycle where each 6 h dark period aligned with the continuous 6 h period when the exemplar wind farm produced the least energy (Figure 1A, 1E). In both cases, dark periods beginning in the final 6 h of a 24 h period continued into the next day..

**Figure 1.**
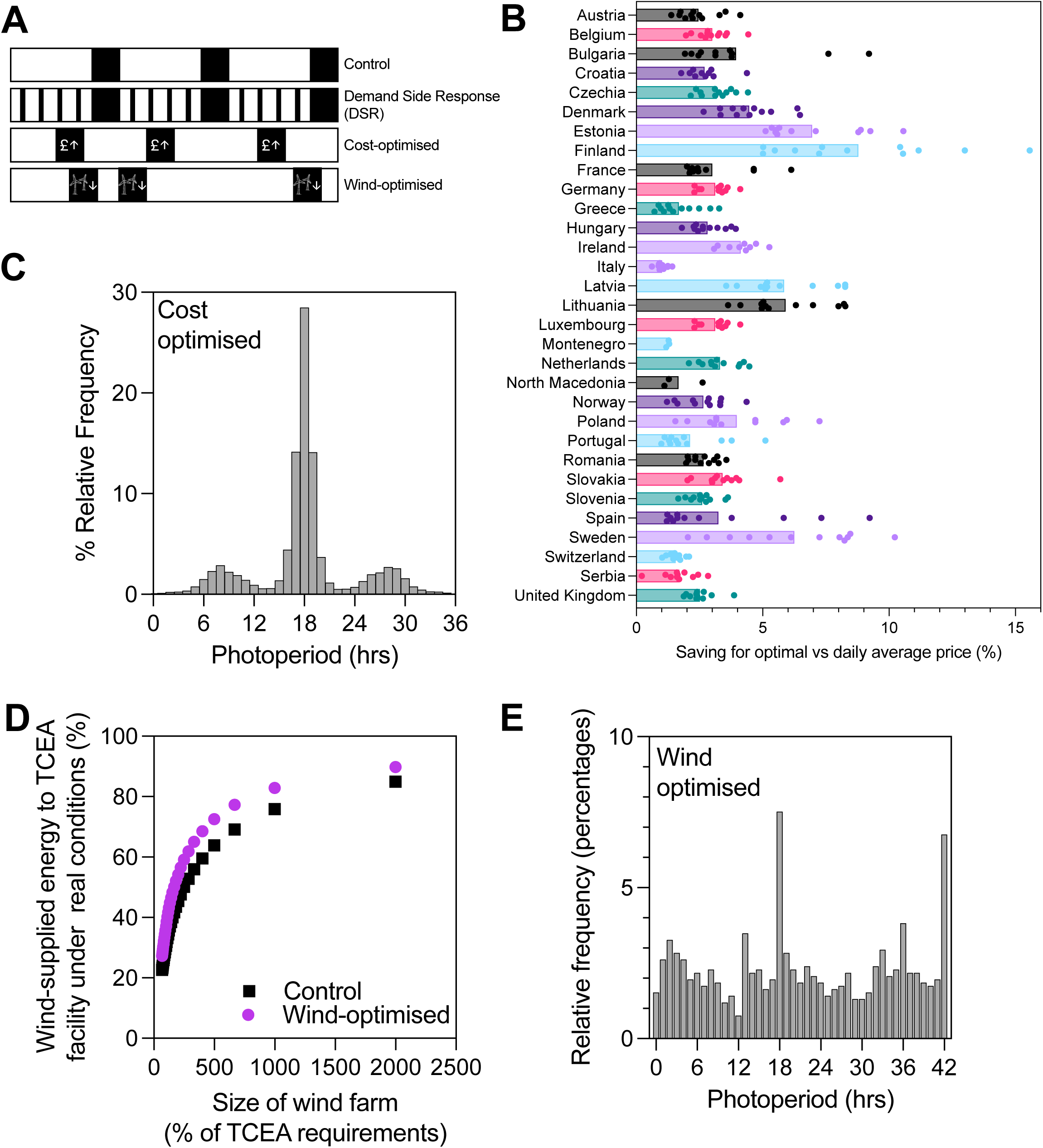
Varied photoperiods can minimize economic and environmental impacts of Total Controlled Environment Agriculture (TCEA). **(A)** Diagram representing an exemplar 72 h under each light regime used in this study. Control conditions comprised a fixed cycle of L18:D6 cycles, alongside Demand Side Response (DSR; consistent L18:D6 cycles with 15 minute dark intervals every 4 h), Cost-optimised (L18:D6 every 24 h with the placement of 6 h night-time corresponding to highest energy pricing within the 24 h day), and wind-optimised (L18:D6 every 24 h but the timing of the 6 hour dark period is adjusted to occur when wind speeds are forecasted to be lowest in the given 24 h). White spaces indicate lit photoperiods, black spaces indicate night-time, dashed blue lines indicate the boundaries of a 24 h period which must contain a 6h night. **(B)** Projected cost savings in different European electricity markets over a 10-year period when implementing our cost-optimised algorithm (2015-2025). See also Supplemental Figure 1. **(C)** Distribution of photoperiod length when aligning TCEA lighting with our cost-optimal algorithm. **(D)** Comparison of windfarm capacity required to supply a proportion of TCEA requirements under regular photoperiods or during application of our wind-optimal algorithm. **(E)** Distribution of photoperiod length when aligning TCEA lighting with our wind-optimised algorithm.

### Phenotypic Characterisation in TCEA Growth Environments

Pilot plant-scale experiments were completed at the University of Glasgow using an IGS Research Rig. Plants were grown with 10-12 trays per growth slot and a mixed population of different crop varieties. Each crop variety was grown on a separate tray and randomly distributed within the growth slot. Seeds were sown onto rehydrated peat-free coconut coir pellets (Jiffy 7-C Coir Pellets, Jiffy Group, Zwijndrecht, Zuid-Holland, Netherlands) in a 40-cell tray (QP D 40/4,5 R, Herkuplast, Germany). Seeds were germinated for 3 days in darkness at 24°C and 70% relative humidity before light regimes began. All plants were grown for 4 weeks (24-25 days) under control, DSR, cost-optimised or wind-optimised light regimes prior to harvest. For each tray, heights were measured for 5 plants per tray taken from the four corners and the centre of the tray before the plants were harvested at the base of stem and fresh weight was measured for each tray-cell individually. Plants were grown under ambient CO_2_ concentrations, with experimental lighting applied in 3 lighting phases (Table 1).

**Table 1.** Lighting qualities for each phase of lettuce grown at IGS and all plants grown at UofG.

| **Phase** | **Duration (days)** | **PPFD (µmol⍰m<sup>-2</sup>⍰s<sup>-1</sup>)** | **Red:Green ratio** | **Red:Blue ratio** | **Blue:Green ratio** | **Red:Far-red ratio** |
|---|---|---|---|---|---|---|
| 1 | 10 | 257 | 23.13 | 2.89 | 8.00 | 20.56 |
| 2 | 8 | 260 | 23.13 | 2.76 | 8.38 | 20.56 |
| 3 | 1 | 264 | 17.90 | 2.39 | 7.50 | 22.38 |

Commercial scale TCEA trials were carried out in the growth towers of Intelligent Growth Solutions Limited (IGS) (https://www.intelligentgrowthsolutions.com/) at the Crop Research Centre in Invergowrie, Dundee, UK. Two seeds were sown in 40-cell trays (QP D 40/4,5 R, Herkuplast, Germany) by a Mosa Drum Seeder sowing machine (Mechanical Botanical, Chiddingfold, United Kingdom) on hydrated peat free coconut coir supplemented by 2% V/V vermiculite (Plant!t, United Kingdom) (Jiffy Indoor Farming coir, Jiffy Group, Zwijndrecht, the Netherlands) and stored in the dark (24°C, 90% relative humidity) to germinate for 2 and 3 days for lettuce and basil, respectively. Once germinated, seedling trays were placed on a liner in the large Growth Tray with Lights (GTL) in a growth tower slots at 24°C and 70% relative humidity under the designated lighting regimes. All plants were watered regularly using standard nutrient solution (NPK molar ratio 5:1:4, EC 2.0, pH 5.8). . Plants were grown at a controlled CO_2_ concentration of 800 μmol mol^−1^. Timing of light-dark periods were based upon experimental conditions, with daily light qualities and intensities during illuminated periods varying with growth phase (Pičmanová et al., 2026). Lettuce plants at IGS were grown under 3 lighting phases (Table 1). Basil plants at IGS were grown under 5 lighting phases (Table 2). Plants in the IGS growth towers were grown in 35 propagation trays per GTLs in monoculture (Supplementary Figure 3). Basil was grown in the IGS growth towers for 20 days and lettuce for 19 days under light. At harvest, 12 propagation trays were selected from each growth population of 35 trays. Heights were measured for 5 plants per tray taken from the four corners and the centre of the tray before the plants were harvested at the base of stem and fresh weight was measured for the whole tray. For dry weights , individual plants were taken at random from the growth population and harvested at the base of stem. Fresh weight was measured before the plants were dried in an oven at 50-70°C for 4-14 days and measured again for dry weights. Harvest data were plotted using GraphPad Prism v.11.0.2.

**Table 2.**
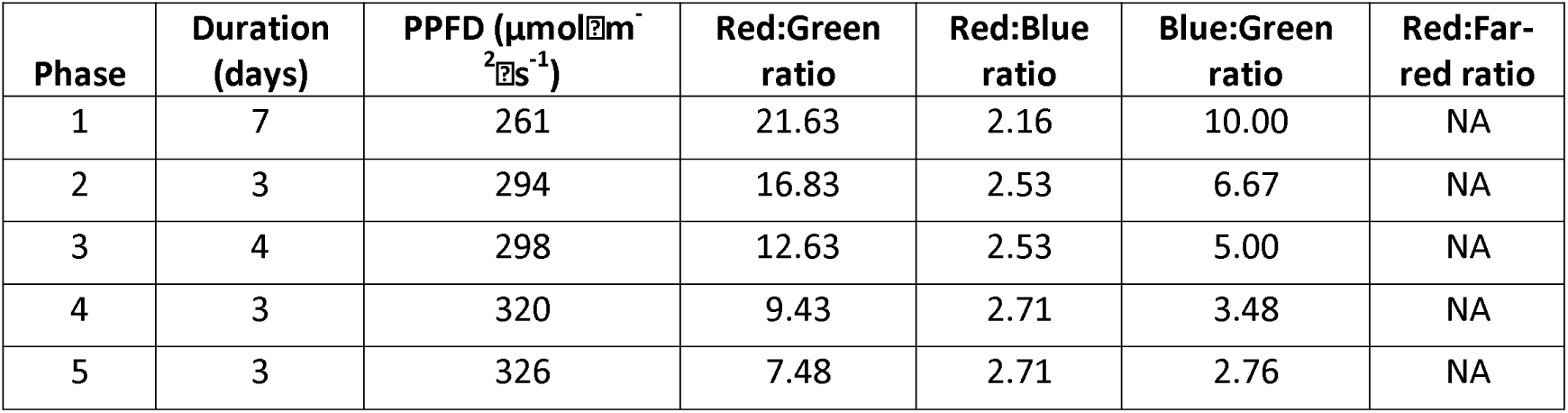
Lighting qualities for each phase for basil grown at IGS.

| **Phase** | **Duration (days)** | **PPFD (µmolm<sup>-2</sup>s<sup>-1</sup>)** | **Red:Green ratio** | **Red:Blue ratio** | **Blue:Green ratio** | **Red:Far-red ratio** |
|---|---|---|---|---|---|---|
| 1 | 7 | 261 | 21.63 | 2.16 | 10.00 | NA |
| 2 | 3 | 294 | 16.83 | 2.53 | 6.67 | NA |
| 3 | 4 | 298 | 12.63 | 2.53 | 5.00 | NA |
| 4 | 3 | 320 | 9.43 | 2.71 | 3.48 | NA |
| 5 | 3 | 326 | 7.48 | 2.71 | 2.76 | NA |

## Results

### Manipulation of lighting regimes refines energy usage to align with economic and environmental priorities

The addition of renewable energy sources to energy distribution grids creates opportunities to align energy usage with availability (Figure 1). Demand-Side Response (DSR) provides opportunities to stabilise the grid, avoiding brown-outs or conversely avoiding grid overload when excess energy is available (Figure 1A). Beyond these short-term interventions, the dynamic pricing presented within modern electricity distribution systems enable further optimisation. For example, the UK’s wholesale electricity market varies on an hourly basis and is priced 24 h in advance. We wanted to understand how these dynamics could be exploited to minimise the cost of energy powering a TCEA facility by aligning night-time with the highest cost periods (Figure 1A, 1B). An analysis of EU countries over 10 years reveals average annual savings of approximately 3% can be achieved, with savings of up to 8% in some markets (Finland, Figure 1B). Savings have become more pronounced in recent years, with recent increases in energy costs driving potential cost reductions in Finland of 15% in 2025 (Supplemental Figure 1). We next assessed how daylength was affected by aligning night-time with high-cost periods (Figure 1C, Supplemental Figure 2). While 18 h photoperiods were the modal value, approximately 10% of days were substantially shorter (∼8 h) or longer (∼28 h), representing periods when the night period was shifted by 10 h to achieve maximal savings. These data suggest that substantial savings can be made with minimal effects upon TCEA operation.

Beyond economic considerations, co-siting TCEA facilities alongside renewable energy sources such as windfarms provides an opportunity to reduce energy draw from the distribution grid, reducing CO_2_ generation and increasing opportunities to diversify the community benefits of the green transition (Figure 1D-1E). We first examined the relationship between windfarm size and the proportion of wind-supplied energy used by the TCEA facility (Figure 1D). A windfarm with a capacity 20x larger than vertical farm maximum load will fulfil 85.0% of energy requirements although smaller windfarm installations are likely to provide a greater cost:benefit ratio given the capital costs of construction (Figure 1D). For example, a wind farm five times the size of the TCEA facility will provide 63.9% of the electricity required, whereas a windfarm four times the size will provide 59.5%. We next examined whether aligning night-time with daily periods when wind strength is weakest would affect the proportion of energy provided to the TCEA facility (Figure 1A, 1D). Wind-optimised light regimes increase the proportion of electricity that can be supplied to a TCEA facility, with a windfarm capacity 2.5 times larger than maximum consumption providing 59.1% of required energy (Figure 1D). Optimising for windfarm generation provides a much more extensive distribution of photoperiod length, with only 8% of lit periods being 18 h (Figure 1E). These data illustrate the distinct consequences of optimising light regimes for economic or environmental priorities.

### Circadian timing buffers the effects of optimised lighting regimes

We were interested how our optimised light regimes affected biological timing and so examined patterns of *CCA1::LUC2* bioluminescence using transgenic Arabidopsis plants (Figure 2; (Battle et al., 2024)). Since lighting is the primary contributor to operational costs (Kozai *et al*. 2016) we implemented DSR by introducing a 15 min interval of darkness every 4 h during the day (Figure 1A, 2D). Wild type Arabidopsis plants maintained a circadian period of 23.82±0.04 h under control conditions which was comparable to that of wild type plants transferred to DSR conditions (23.83±0.01 h; Figure 2A-2C). However, despite the maintenance of period, circadian phase was delayed by 1.5 h in wild-type Arabidopsis during implementation of DSR suggesting the circadian system does respond to the application of demand flexing conditions (Figures 2A, 2D).

**Figure 2.**
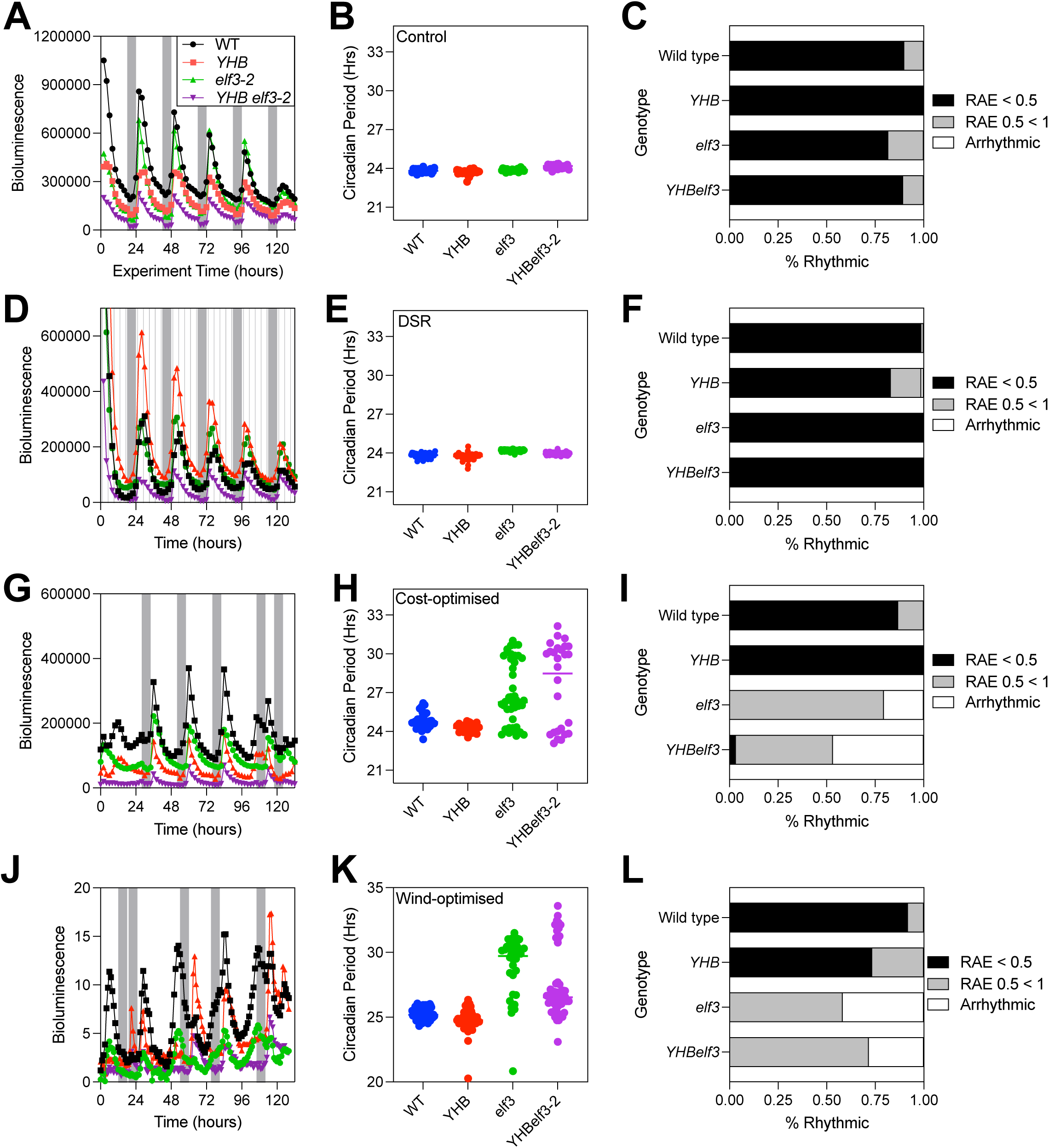
Circadian Rhythms are Maintained in Arabidopsis Plants Grown Under Energy Use Demand Management. **(A)** Bioluminescence derived from *CCA1::LUC2* expression in plants grown under control L18:D6 cycles. **(B)** Circadian period of plants grown under driven L18:D6 cycles in (A). **(C)** Proportion of plants that retained bioluminescence rhythms under control L18:D6 cycles. **(D)** Bioluminescence derived from *CCA1::LUC2* expression in plants grown under Demand Side Response (DSR) regimes. **(E)** Circadian period of plants grown under driven DSR cycles in (D). **(F)** Proportion of plants that retained bioluminescence rhythms under DSR cycles. **(G)** Bioluminescence derived from *CCA1::LUC2* expression in plants grown under cost-optimised lighting regimes. **(H)** Circadian period of plants grown under cost-optimised regimes in (G). **(I)** Proportion of plants that retained bioluminescence rhythms under cost-optimised cycles. **(J)** Bioluminescence derived from *CCA1::LUC2* expression in plants grown under wind-optimised lighting regimes. **(K)** Circadian period of plants grown under wind-optimised regimes in (J). **(L)** Proportion of plants that retained bioluminescence rhythms under wind-optimised cycles. Rhythms were considered robust if circadian period was estimated within 16 and 30h, with a Relative Amplitude Error (RAE) < 0.5. Plants returning poor rhythms (RAE > 0.5) and those that were arrhythmic are also reported.

We next assessed the contribution of circadian timing to the driven patterns of *CCA1::LUC2* observed (Figure 2A-2C). *elf3* plants cannot maintain circadian rhythms whereas *YHB* seedlings express a constitutively active phyB photoreceptor and are consequently less responsive to light (Hu et al., 2009; Su and Lagarias, 2007; Hu et al., 2020). The combined *YHBelf3* double mutant has limited phenotypic plasticity and lacks circadian rhythms (Battle et al., 2024). Since circadian timing modulates the responses of plants to environmental signals, we expected *elf3* seedlings to respond uniformly to dawn regardless of photoperiod length whereas *YHB* seedlings would be less responsive to irradiation (Figure 2). As previously reported, regular light:dark cycles were sufficient to drive rhythmic patterns of *CCA1::LUC2* bioluminescence in otherwise arrhythmic *elf3* and *YHBelf3* seedlings (Figure 2A-2C); (Battle et al., 2024). *elf3* and *YHBelf3* plants retained rhythms in DSR conditions, and did not induce *CCA1::LUC2* expression after a 15 min dark interval (Figure 2D). However, we were interested to note that the phase of *CCA1::LUC2* bioluminescence was not delayed in *YHBelf3* plants in DSR conditions compared to wild type (Figure 2D-2F).

In cost-optimised conditions, wild-type and *YHB* plants were able to retain circadian rhythms of CCA1-driven bioluminescence despite dawn induction of CCA1::LUC2, consistent with behaviour previously observed for CCA1 RNA (Figure 2G-2H, (Wang et al., 1997)). Circadian gating was evident in the magnitude of bioluminescence induced in wild-type and *YHB* in response to dawn, with extended daylengths also provoking premature anticipation of dawn in both genotypes (Figure 2G). By contrast, patterns of elf3 and YHBelf3 bioluminescence were primarily driven by dawn, with no anticipatory or gating behaviour or observed (Figure 2G). The reduced circadian function and responsiveness of elf3 and YHBelf3 seedlings to dawn led to a loss of detectable rhythmicity in elf3 and YHBelf3 plants transferred to cost-optimised conditions (Figure 2H+2I).

Although utilising more varied daylengths, wind-optimised regimes elicited similar dawn responses and anticipatory behaviours of *CCA1::LUC2* expression (Figures 1C, 1E, 2J). Circadian period was extended in wildtype plants although 92% of wild type maintained a circadian rhythm in wind-optimised conditions (25.26±0.44h; Figure 2K+2L). Wild-type and *YHB* seedlings continued to maintain a degree of rhythmicity despite the irregular daylengths (Figure 2J-L). By contrast patterns of *CCA1::LUC2* bioluminescence were completely irregular and lost all rhythms in *elf3* and *YHBelf3* seedlings, with dawn being the primary driver of varied bioluminescence in these lines (Figure 2K+L). Indeed, both *elf3* and *YHBelf3* seedlings increased *CCA1::LUC2* bioluminescence after a single hour of irradiation (Figure 2J). Our data suggest that circadian rhythmicity is maintained in wild type plants in cost- and wind-optimised conditions despite the clock retaining responses to day/night transitions. Genetic disruption of the circadian system is sufficient to abrogate these effects.

### Optimised Light Regimes Affect Arabidopsis Growth

Circadian resonance has been proposed as a mechanism to promote crop yield by aligning circadian period with daylength and so we next examined whether our optimised lighting regimes affected growth and development (Dodd et al., 2005; Anton-Sales et al., 2026). Wild type *Arabidopsis thaliana* plants grown under DSR conditions accumulated equivalent biomass as those grown in regular 18:6 days although flowering time was delayed by approximately two days, possibly due to the modest reduction in daylength or amount of light provided (Figures 1A, 3A-3D). We next grew plants under cost-optimised or wind-optimised regimes and were interested to note that Arabidopsis plants accumulated less dry weight in these regimes compared to control conditions despite the treatments supplying the same total number of photons during the experiment (p < 0.05, Figure 3B). Rosette area was unaffected by the applied light regimes, suggesting differences in leaf thickness contribute to dry weight differences (Figure 3C). Growing wild type Arabidopsis plants under cost-optimised conditions did not affect flowering time (p > 0.05, Figure 3D) although wildtype plants grown in a wind-optimised regime flowered earlier than those in control conditions (p < 0.05, Figure 3D). These data demonstrate the sensitivity of Arabidopsis plants to varying light regimes, with different aspects of their life history being affected dependent upon the regime applied.

**Figure 3.**
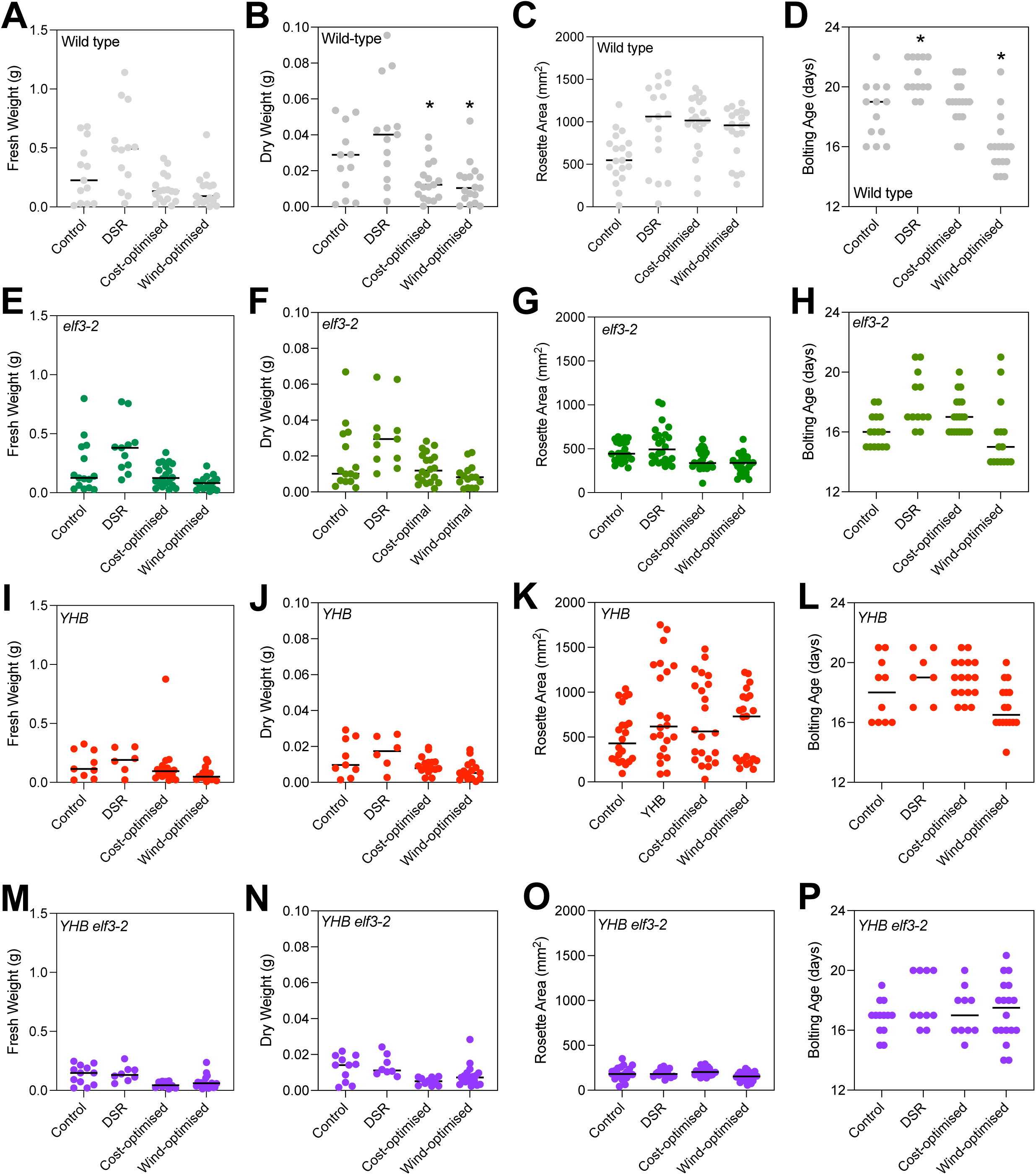
Life Cycle Characterisation of Arabidopsis plants grown under Energy Use Demand Management. (A,E,I,M) Fresh weight harvest of *Arabidopsis thaliana* plants grown under different light regimes. Data from wild type (A), *elf3-2* (E), *YHB* (I), and *YHBelf3-2* (M) plants are presented. **(B, F, J, N)** Dry weight harvest of *Arabidopsis thaliana* plants grown under different light regimes. Data from wild type (B), *elf3-2* (F), *YHB* (J), and *YHBelf3-2* (N) plants are presented. **(C, G, K, O)** Rosette surface area of plants at bolting. Data from wild type (C), *elf3-2* (G), *YHB* (K), and *YHBelf3-2* (O) plants are presented. **(D, H, L, P)** Age of plants at bolting under different lighting regimes. Data from wild type (D), *elf3-2* (H), *YHB* (L), and *YHBelf3-2* (P) plants are presented. Plant biomass and rosette area was measured immediately after bolting. Significant comparisons are presented from a two-way ANOVA, adjusted using Tukey’s multiple-comparisons test ( \**p*=0.01–0.05).

We next assessed whether disruption of circadian timing would be sufficient to rescue the impaired growth observed (Figure 3E-3H). We were interested to note that *elf3* plants did not display a yield penalty when grown in DSR, cost-optimised or wind-optimised conditions (Figure 3E-3H; (Hicks et al., 1996; Thines and Harmon, 2010). However, as previously reported, *elf3* plants have reduced vegetative growth in control conditions compared to wild type plants (Figure 3A, 3E; (Battle et al., 2024)). The observed reduction in vegetative growth is likely related to the earlier flowering time observed in *elf3* plants (Figure 3D, 3H; (Zagotta et al., 1996; Battle et al., 2024)). *elf3* flowering time was consistently earlier than wildtype, with this flowering phenotype unaffected by the applied lighting regime (Figure 3D, 3H). These data suggest that disruption of circadian timing can mitigate the effects of irregular daylengths upon plant growth. However, the pleiotropic effects of the *elf3* mutation upon circadian timing, photomorphogenesis, and flowering time complicate data interpretation and limit the commercial applicability of the phenotype (Figure 3E-3H, (Battle et al., 2024)).

The photomorphogenic consequences of *YHB* expression are epistatic to *elf3*, with growth of *YHBelf3* plants indistinguishable from *YHB* seedlings in control conditions (Figure 3I-3P; Battle et al., 2024). *YHB* plants and *YHBelf3* grew consistently regardless of the lighting regime implemented (Figure 3I-3K). Similarly, flowering time was unchanged in *YHB* and *YHBelf3* seedlings when grown in each of the optimised conditions (Figure 3L and 3P). These data demonstrate that genetic manipulation of light signalling and circadian pathways can reduce phenotypic plasticity in response to environmental cues, providing a route to developing consistently-performing crops when required.

### End User Energy Demand management provides opportunities for commercial-scale growth

We wanted to examine how crop growth was affected by demand side flexibility so that growers can understand the trade-offs associated with participating in these different schemes (Figure 4). We first examined growth of six different crops in our DSR conditions in our IGS Research RIG. We observed a significant increase in basil fresh weight (p < 0.05, 2-way ANOVA), although there was no significant yield difference in Green Batavia lettuce, Tango lettuce, mustard greens, or rocket (Figure 4A-4E). Interestingly, there was no significant difference in basil dry weight under DSR conditions suggesting that the observed differences were due to water content rather than biomass accumulation (Figure 4F). By contrast, growing crops under cost-optimised cycles had a significant effect upon the harvested fresh weight yield of mustard greens (falling by 32%; Figure 4D, p < 0.05) but not the other crops examined. As in DSR conditions, this disparity appeared to be due to water content as no difference observed in dry weight of mustard greens harvested from control or cost-optimised conditions (Figure 4I). Wind-optimised regimes similarly had minimal effect upon fresh weight accumulation compared to controls in any of the crops tested (Figure 4). Rocket displayed a 10% increase in harvested fresh weight, which was also attributed to increased water content as there was no significant difference in dry weight (Figure 4E, p < 0.05, 2-way ANOVA). Interestingly, dry weight harvests of both Green Batavia and Tango lettuce were increased in wind-optimised conditions despite there being no difference in fresh weight (Figure 4G, 4H, p < 0.05, 2-way ANOVA). These data suggest that varied photoperiods can be implemented in TCEA without impairing crop growth.

**Figure 4.**
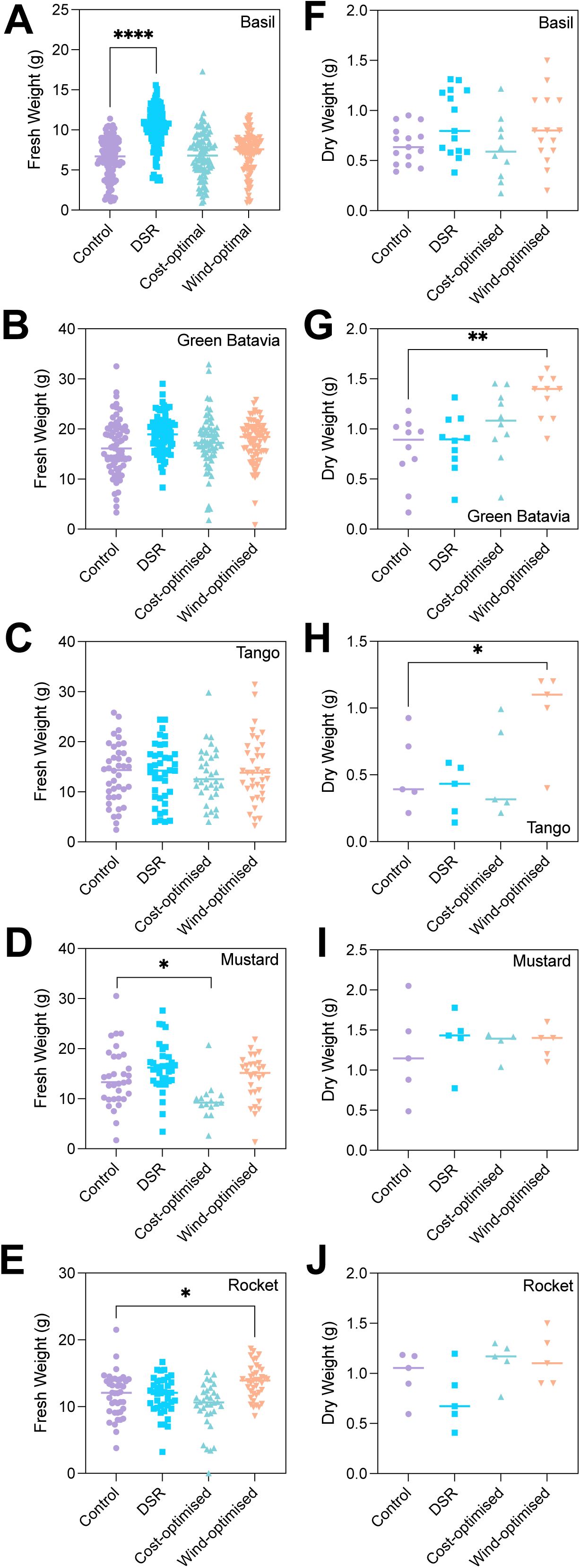
Harvest Comparison of Crops Grown Under Energy Use Demand Management at Pilot Plant Scale. (A, **F)** Fresh weight (A) and dry weight (F) of basil plants grown under different lighting regimes. **(B, G)** Fresh weight (B) and dry weight (G) of Green Batavia lettuce plants grown under different lighting regimes. **(C, H)** Fresh weight (C) and dry weight (H) of Tango lettuce plants grown under different lighting regimes. **(D, I)** Fresh weight (D) and dry weight (I) of mustard plants grown under different lighting regimes. **(E, J)** Fresh weight (E) and dry weight (J) of mustard plants grown under different lighting regimes. Plants were grown for 24 days under 18L:6D fixed cycle control or energy use demand management regimes. Significant comparisons are presented from a two-way ANOVA, adjusted using Tukey’s multiple-comparisons test ( * *p* < 0.05, ** *p* <0.01, **** *p* <0.0001).

We next assessed growth of a subset of our crop panel in a larger commercial scale growth facility (IGS growth towers; Figure 5). Application of a DSR regime had no effect upon the fresh weight of basil plants although there was an 18% decrease in harvested dry weight (Figure 5A and 5B). Basil plants were also shorter in DSR conditions (Figure 5C). By contrast, neither cost-nor wind-optimised conditions affected growth of basil when grown at commercial scale. Harvested fresh weight of Green Batavia lettuce was reduced under DSR conditions although dry weight was comparable to control conditions (Figure 5D and 5E). As for basil, fresh weight of Green Batavia lettuce was unaffected by cost-optimal or wind-optimal conditions although lettuce plants were shorter under wind-optimal lighting (Figure 5D-5F).

**Figure 5.**
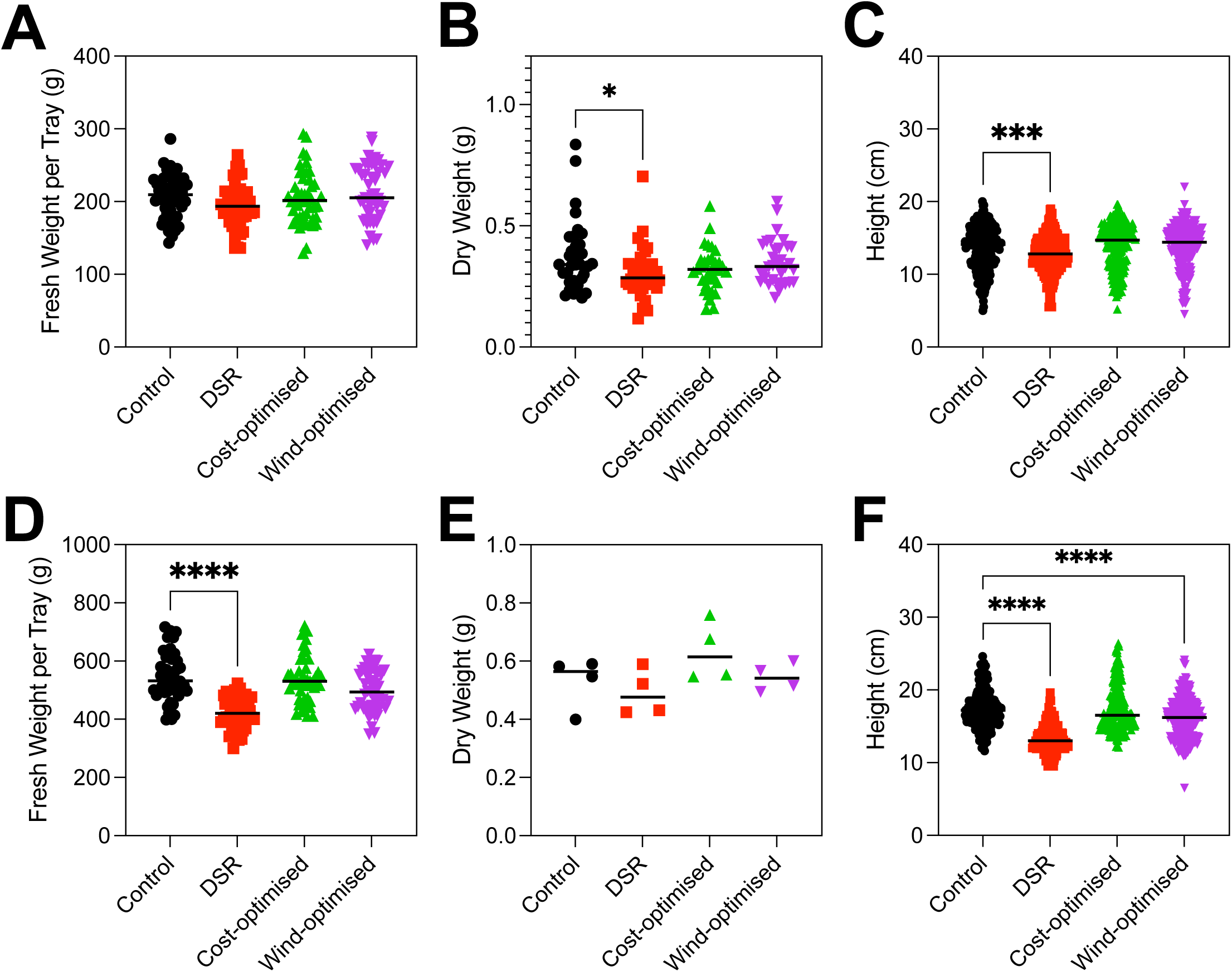
Harvest Comparison of Crops Grown Under Energy Use Demand Management at Commercial Scale. (A, **B)** Fresh weight (A) and dry weight (B) of basil plants grown under different lighting regimes. **(C)** Height of basil plants grown under different lighting regimes. **(D, E)** Fresh weight (D) and dry weight (E) of Green Batavia plants grown under different lighting regimes. **(F)** Height of Green Batavia lettuce plants grown under different lighting regimes. Plants were grown for 19 and 20 days (lettuce and basil, respectively) under 18L:6D fixed cycle control or energy use demand management regimes. Significant comparisons are presented from a two-way ANOVA, adjusted using Tukey’s multiple-comparisons test (* *p* < 0.05, *** *p* <0.001, **** *p* <0.0001).

## Discussion

### Demand Side Management provides opportunities to reduce energy expenditure in TCEA

The rapid expansion of renewable electricity generation is increasing the value of flexible electricity demand across modern energy systems (Fu et al., 2024). Renewable installations are frequently constrained by grid capacity and transmission infrastructure, particularly in remote regions where wind resources are strongest (Tröndle et al., 2020). Co-locating flexible agricultural loads with renewable generators therefore provides an additional route to increase utilisation of locally generated electricity while creating new economic opportunities for rural communities (Kaiser et al., 2024; Shah et al., 2023). Our study demonstrates that Total Controlled Environment Agriculture (TCEA) can provide meaningful demand-side flexibility while maintaining commercially acceptable crop performance under a range of operational scenarios (Figure 1 and 5).

Each of the end user energy demand scenarios implemented had different consequences for the growth environment (Figure 1). Demand-Side Responses (DSR) implemented through 15 min dark intervals had relatively minor effects on daylength, whereas the cost-optimised and wind-optimised schedules imposed substantially greater variation (Figure 1A, 1C, 1E). We chose to implement a ‘high-use’ DSR regime to understand how short-term lighting reductions affect plant growth, although data suggests that DSR is implemented much less frequently in practice (NESO 2026). Despite our DSR regime reducing lighting by 8%, yield was maintained relative to control conditions in the majority of crops and facilities examined (Figures 3, 4, 5). Since DSR contracts have financial incentives, it is likely that commercial TCEA facilities will benefit from the implementation of these approaches. Beyond these short-term, responsive light regimes, analyses of historical electricity pricing suggest that aligning photoperiods with periods of lower electricity cost could reduce operating expenditure by approximately 3% across European markets, with substantially greater savings in specific regions and years (Figure 1). Alternately, aligning lighting schedules with periods of high wind generation provides the opportunity to increase the proportion of energy supplied directly from co-located renewable infrastructure (Figure 1). Together, these findings establish TCEA as a potential participant in future smart energy networks, delivering both agricultural outputs and grid-balancing services.

### Crop productivity can be maintained during irregular photoperiods

The natural light environment is complex and heterogenous with daily changes in cloud cover coinciding with longer term, seasonal patterns of photoperiod and context-dependent shading. Plants have consequently evolved numerous adaptations to maximise fecundity, ranging from rapid changes in gene expression through to longer term phenotypic plasticity encompassing varied vegetative growth and flowering time (Jones, 2018). TCEA can exploit this phenotypic plasticity to control traits underlying yield and quality, although phenotypic plasticity could conversely reduce yield or quality when aligning lighting regimes with economic and/or environmental considerations.

Indoor crops tend to be grown under lower light levels than in the field and are consequently source-limited; meta-analysis demonstrates that increasing light elicits a corresponding benefit in crop biomass (Kaiser et al., 2024). Despite this sensitivity, aligning fluence rate with energy cost during the photoperiod does not affect fresh weight harvest in many crops when total irradiance is maintained (Mäkinen et al., 2025; Kaiser et al., 2024). Our work extends these findings to crops grown under irregular photoperiods, demonstrating immediate economic and environmental benefits for TCEA when dark periods coincide with periods of high energy cost or reduced energy availability across the grid (Figures 4 and 5).

The circadian clock coordinates daily cellular behaviour with environmental signals, with increased yield reported when circadian period is aligned with daylength (Anton-Sales et al., 2026; Dodd et al., 2005). Disruption of the circadian system using the *elf3* allele reduced vegetative growth in all conditions tested, likely due to the previously reported early flowering phenotype (Zagotta et al., 1996; Battle et al., 2024). The developmental consequences of circadian disruption therefore appeared epistatic to the imposition of irregular photoperiods, although there was no difference in the fresh- or dry-weight of *elf3* plants grown under our different conditions that could be interpreted as a loss of circadian resonance (Figure 3; (Anton-Sales et al., 2026)).

Since the circadian system is capable of rapid entrainment to different photoperiods we hypothesised that the highly variable daylengths generated by cost- and wind-optimised regimes would disrupt circadian timing (Sanchez et al., 2020). Instead, although DSR delayed circadian phase and wind-optimised regimes lengthened circadian period, the majority of wild-type Arabidopsis plants retained detectable rhythms demonstrating the considerable robustness of the circadian system (Figure 2). These data suggest that multiple days of entrainment are required to align plant circadian rhythms with changing environmental signals despite rapid induction of transcripts such as CCA1 at dawn (Figure 2G, 2J). It is likely that metabolic daylength measurement, photoreceptors, and retrograde signalling pathways all contribute to the maintenance of circadian timing in these irregular conditions (Harmer, 2025; Li and Gendron, 2025; Jones, 2019; Sanchez et al., 2020).

## Conclusions and future directions

The chronoculture concept highlights how treatments at specific time of day can promote desirable traits and minimise costs associated with fertiliser and herbicide treatment (Ogasawara et al., 2025; Steed et al., 2021). Our data suggest the chronoculture concept can additionally exploit endogenous circadian rhythms to alleviate the effects of variable photoperiods (Figures 2 and 3). Importantly, the retention of circadian rhythms under our cost- and wind-optimised regimes suggest that chronoculture approaches remain applicable when applying end user energy demand management (Ogasawara et al., 2025; Steed et al., 2021). Varying light irradiation during the day in addition to altering photoperiod will likely provide additional economic and environmental benefits to commercial growers.

---

**Supplemental Figure 1.** Projected cost savings in European electricity markets over a 10-year period when implementing our cost-optimised algorithm (2015-2025).

**Supplemental Figure 2.** Distribution of photoperiod length when aligning TCEA lighting with our cost-optimal algorithm in different European energy markets (2015-2025).

