## Supplemental Figure 1 for "Harnessing Circadian Resilience for Demand-Flexible Controlled Environment Agriculture"

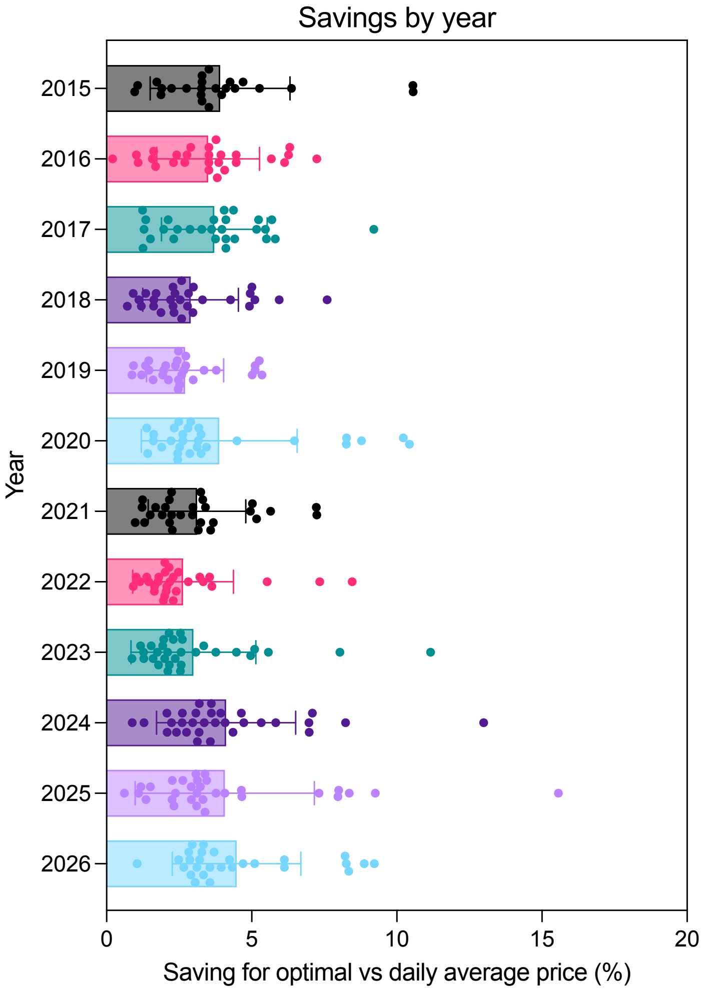


**Supplemental Figure 1.** Projected cost savings in European electricity markets over a 10-year period when implementing our cost-optimised algorithm (2015-2025).
