## Supplemental Figure 2 for "Harnessing Circadian Resilience for Demand-Flexible Controlled Environment Agriculture"

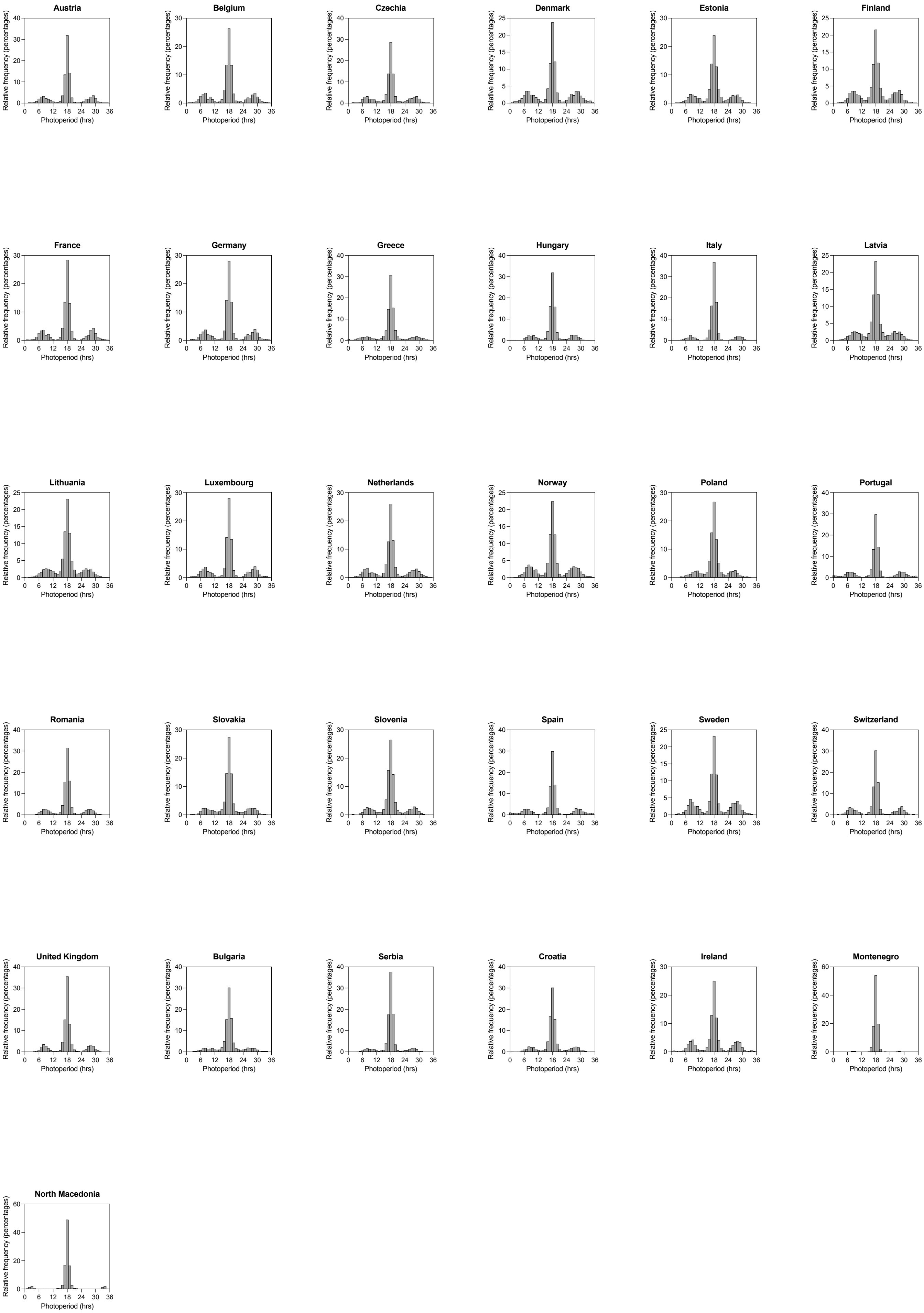


**Supplemental Figure 2.** Distribution of photoperiod length when aligning TCEA lighting with our cost-optimal algorithm in different European energy markets (2015-2025).
